# Sequence amplification via cell passaging creates spurious signals of positive adaptation in influenza H3N2 hemagglutinin

**DOI:** 10.1101/038364

**Authors:** Claire D. McWhite, Austin G. Meyer, Claus O. Wilke

## Abstract

Clinical influenza A isolates are frequently not sequenced directly. Instead, a majority of these isolates (~70% in 2015) are first subjected to passaging for amplification, most commonly in non-human cell culture. Here, we find that this passaging leaves distinct signals of adaptation in the viral sequences, which can confound evolutionary analyses of the viral sequences. We find distinct patterns of adaptation to Madin-Darby (MDCK) and monkey cell culture absent from unpassaged hemagglutinin sequences. These patterns also dominate pooled datasets not separated by passaging type, and they increase in proportion to the number of passages performed. By contrast, MDCK-SIAT1 passaged sequences seem mostly (but not entirely) free of passaging adaptations. Contrary to previous studies, we find that using only internal branches of the influenza phylogenetic trees is insufficient to correct for passaging artifacts. These artifacts can only be safely avoided by excluding passaged sequences entirely from subsequent analysis. We conclude that future influenza evolutionary analyses should appropriately control for potentially confounding effects of passaging adaptations.

## 1. Introduction

The routine sequencing of clinical isolates has become a critical component of global seasonal influenza surveillance (World Health Organization Global influenza surveillance network, 2011). Analysis of these viral sequences informs the selection of future vaccine strains(Stöhr et al., 2012; WHO Writing Group et al., 2012), and a wide variety of computational methods have been developed to identify sites under selection or immune-escape mutations(Blackburne et al., 2008;, Koelle et al., 2006; Nelson et al., 2006; Suzuki, 2008; Wolf et al., 2006), or to predict the short-term evolutionary future of influenza virus (Łuksza and Lässig, 2014; Neher et al., 2014). However, sites that appear positively selected in sequence analysis frequently do not agree with sites identified experimentally in hemagglutination inhibition assays (Kratsch et al., 2016; Meyer and Wilke, 2015; Tusche et al.,2012), and the origin of this discrepancy is unclear. Here, we argue that a major cause of this discrepancy is widespread passaging of influenza virus before sequencing.

Clinical isolates are often passaged in culture one or more times to amplify viral copy number, as well as to introduce virus into a living system for testing strain features such as vaccine response, antiviral response, and replication efficiency (Kumar and Henrickson, 2012; World Health Organization Global influenza surveillance network, 2011). A variety of culture systems are used for virus amplification. Cell cultures derived from Madin-Darby canine kidney (MDCK) cells are by far the most widely used system, with the majority of sequences in influenza repositories deriving from virus that has been passaged through an MDCK or modified MDCK cell culture (Balish et al., 2005; Bogner et al., 2006). Influenza virus may also be passaged through monkey kidney (RhMK or TMK) cell culture or injected directly into egg amniotes. Alternatively, complete influenza genomes can be obtained from PCR-amplified influenza samples without intermediate passaging (Katz et al., 1990; Lee et al., 2013a).

Several experiments have demonstrated that influenza virus hemagglutinin (HA) accumulates mutations following rounds of passaging in both cell (Ilyushina et al., 2012; Lee et al., 2013b; Wyde et al., 1977) and egg culture (Robertson et al., 1993). The decreased number of mutations in MDCK-based cell culture is the main argument for use of this system over egg amniotes in vaccine production (Katz and Webster, 1989), with MDCK cells expressing human SIAT1 having the highest fidelity to the original sequence and reduced host adaptation (Hamamoto et al., 2013). Viral adaptations to eggs were recently linked to reduced vaccine efficacy (Skowronski et al., 2014; Xie et al., 2015) and were implicated as potentially contributing to reduced efficacy of 2014-2015 seasonal H3N2 influenza vaccination in the World Health Organization’s recommendations for 2015-2016 vaccine strains (The World Health Organization,2015). As the majority of influenza vaccines worldwide are produced in eggs, vaccine strain selection is limited to virus with the ability to replicate rapidly in this system (World Health Organization Global influenza surveillance network, 2011).

Although egg-passaged sequences are increasingly excluded from influenza virus phylogenetic analysis (see e.g. the NextFlu tracker (Neher and Bedford, 2015)), due to the known high host-specific substitution rates, cell culture is generally not thought to be sufficiently selective to produce a discernable evolutionary signal. One of few existing evolutionary analyses of passaging effects on influenza virus (Bush et al., 2000) found that passaging caused no major changes in clade structure between egg and cell passaged viruses. However, several studies have recommended the use of internal branches in the phylogenetic tree to reduce passaging effects in evolutionary analysis of Influenza A (Bush et al., 2001; Suzuki, 2006). Another study discovered egg culture to be the cause of misidentification of several sites under positive selection in Influenza B (Gatherer, 2010), but this study was limited to comparing egg-cultured to cell-cultured virus. As the availability of unpassaged influenza sequences has dramatically increased over the past ten years, we can now perform a direct comparison of passaged to circulating virus.

Here, we compare patterns of adaptation in North American seasonal H3N2 influenza HA sequences derived from passaged and unpassaged virus. We divide viral sequences by their passaging history, distinguishing between unpassaged clinical samples, egg amniotes, RhMK (monkey) cell culture, and generic/MDCK-based cell culture. For the latter, we also distinguish between virus passaged in MDCK-SIAT1 cell culture (SIAT1) and in unmodified MDCK or unspecified cell culture (non-SIAT1). We find clear signals of adaptation to the various passaging conditions, and demonstrate that passaging artifacts become more severe with additional rounds of passaging. These signals are strongly present in the tip branches of the phylogenetic trees but can also be detected in internal and trunk branches. We demonstrate the accumulation of these passaging artifacts with additional rounds of serial passaging in non-SIAT1 cells.

Finally, we demonstrate that the identification of antigenic escape sites from sequence data has been confounded by passaging adaptations, and that the exclusion of passaged sequences allows us to use sequence and structural data to highlight regions involved in antigenic escape.

## 2. METHODS

### 2.1 Influenza Sequence Data

Non-laboratory strain H3N2 hemagglutinin (HA) sequences collected in North America were downloaded from The Global Initiative for Sharing Avian Influenza Data (GISAID) (Bogner et al., 2006) for the 1968-2015 influenza seasons. We used exclusively North American sequences to reduce regional variation between influenza virus strains. Non-complete HA sequences were excluded. Sequences were trimmed to open reading frames, filtered to remove redundancies, and aligned by translation-alignment-back-translation using MAFFT (Katoh and Standley, 2013) for the alignment step. Sequence headers of FASTA files were standardized into an uppercase text format with non-alphanumeric characters replaced by underscores. As H3N2 strains have experienced no persistent insertion or deletion events, we deleted sequences that introduced gaps to the alignment. To ascertain overall data quality, we built a phylogenetic tree of the entire sequence set (using FastTree 2.0 compiled for short branch lengths (Price et al., 2010)) and checked for any abnormal clades or other unexpected tree features. We found one abnormal clade of approximately 20 sequences with an exceptionally long branch length (> 0.01) and removed the sequences in that clade from further analysis. Our final dataset consisted of 6873 sequences from 2005-2015 as well as one outgroup of 45 sequences from 1968-1977 (not considered for further analysis). We did not consider sequences collected from 1978-2004.

### 2.2 Identification of Passage History and Evolutionary-Rate Calculations

We divided sequences into groups by their passage-history annotation and collection year,determining passage history by parsing with regular expressions for keywords in FASTA headers (Table 1). We classified 1133 sequences with indeterminate or missing passage histories, or passage through multiple categories of hosts (i.e. both egg and cell), as “other”. The final datasets for individual passage groups contained between 79 and 3041 sequences (Table 1).

**Table 1.**
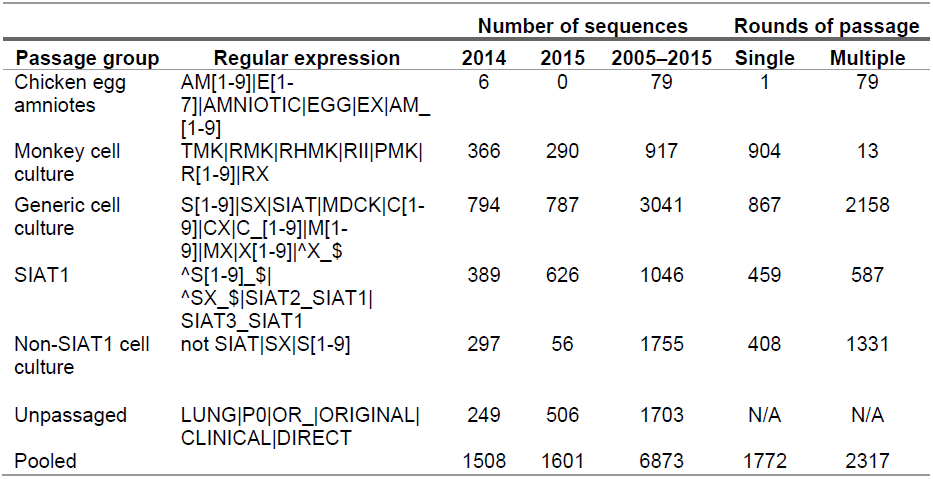
Parsing of passage-annotated FASTA sequences into passage history groups. For each passage group, we defined a regular expression that could reliably identify sequences with that passage history. Regular expressions were applied through the built-in python library “re”. SIAT1 and non-SIAT1 cell culture regular expressions were applied to the subset of sequences identified as generic cell culture sequences. The three middle columns list the number of sequences we identified for each passage group, for years 2014 only, 2015 only, and 2005–2015. The two furthest right columns list the number of single and multiply passaged sequences from 2005–2015 for each condition. Number of sequences Rounds

We additionally divided passage groups into singly and serially passaged subgroups. Sequences matching the regular expression “2|3|4|5|6|1_C|X_|X_C|AND_MV1|X_S|1_S|CX|MX|C1S1|EX|X_E|MIX_RHMK|RII” were classified as having been passaged two or more time.All remaining sequences were passaged only once.

To determine the number of times a sequence was passaged, we used a different set of regular expressions, which we applied only to non-SIAT1 cell-passaged sequences. We first excluded sequences with an indeterminate number of passages by excluding sequences whose record IDs matched the regular expression“^X_$|DETAILS__MDCK|MX_C|^X_C1_|^MX_$|CX_C1|X_C1|DETAILS ND”. We then collected multiply passaged sequences using the regular expression “3|1_C2|2_C1|M1M1_C1|1_MDCK2|4|2_C2|3_C1|1_C3|5|2_C3|3_C2” and doubly passaged sequences using the regular expression "2|1_C1". The remaining sequences were only passaged a single time.

We next constructed phylogenetic trees for each passage group as well as one tree for a pooled dataset combining all individual passage groups and other sequences. All phylogenetic trees were constructed using FastTree 2.0 (Price et al., 2010). We calculated site-specific *dN/dS* values using a one-rate SLAC (Single-Likelihood Ancestor Counting) model implemented in HyPhy (Pond et al., 2005). One rate models fit a site-specific *dN* and a global *dS*, where the global *dS* is the meansite-wise*dS* for a given condition (Spielman et al., 2015). Among different one-rate, site-specific models, SLAC performs nearly identically to other approaches (Spielman et al., 2015), and it was chosen here due to its speed and ease of extracting *dN/dS* estimates along internal and tip branches. To obtain internal and tip branch-specific estimates, we extracted the *dN/dS* values calculated by the SLAC algorithm. We manually calculated*dN/dS* along trunk branches by counting the number of synonymous and non-synonymous substitutions at each trunk site. We defined the trunk as the sequence of branches from the root to the penultimate node before a randomly chosen terminal sequence from the most recent year represented in the tree.

We chose sequences from 2005-2015 as our sample set due the low number of available sequences prior to this period. As *dN/dS* estimates can be confounded by sample size (Spielman et al., 2015), we sought to limit this effect by down-sampling each experimental set to match the number of sequences in the smallest group being considered in a particular analysis (Table 1). To reduce season-to-season variation in the comparison of unpassaged, SIAT1, and non-SIAT1 cell culture, we performed one analysis with sequences from only 2014, which is the year that maximizes sequences available from all three conditions (*n*=249 each).

### 2.3 Geometric Analysis of *dN/dS* Distributions

For each amino acid site *i* in HA, we computed the inverse Euclidean distance to each amino acid site *j*(*j≠i*) in the 3D crystal structure. For each site *i*, we then correlated the list of inverse distances to sites *j* with site-wise *dN/dS* values at sites *j*. This procedure resulted in a correlation coefficient for each site *i*, and we mapped these correlation coefficients onto the corresponding sites *i* in the 3D structure model of HA. In this analysis, sites spatially closest to positively selected regions in the protein yielded the highest correlation coefficients. Thus, this approach allowed us to visualize regions of increased positive selection. As the correlation coefficient for site *i*=224 is consistently highest for sets of sequences undivided by passage history, we chose this site as a reference to compare passage conditions. See Meyer and Wilke (2015) for additional discussion of this approach.

We processed the HA PDB structure to allow for easy alignment with site-wise measures as discussed previously (Meyer and Wilke, 2015). We provided a renumbered and formatted H3N2 HA structure derived from PDBID:2YP7 (2YP7clean.pdb) (Lin et al., 2012) with our data analysis code (see below). Noting that the hemagglutinin protein and gene numbering is offset by 16, all site numbering in this manuscript refers to the protein site position. The alignment of gene and protein numbering schemes to amino acid sequence is available in each supplementarydata file.

### 2.4 Local Branching Index Analysis

We used the framework and code (https://github.com/rneher/FitnessInference) from Neher et al. (2014) to rank sequences according to their Local Branching Index (LBI), a metric that uses branching density to predict progenitor lineages. To build our sample set we divided sequences by year and passage history. We then down-sampled alignments to 70% of the available sequences, up to a maximum of 100 sequences. We repeated each down-sampling fifty times for each condition. We then ranked sequences in each sample according to the LBI algorithm(script rank_sequences.py available from https://github.com/rneher/FitnessInference) and calculated the hamming distance of the top ranked sequence from each conditionto theancestrally reconstructed root sequence of the following year’s unpassaged and pooled trees. The hamming distance derived from the top ranked sequence was divided by the hammingdistance of a randomly chosen sequence from the same condition, to assess if a predicted progenitor was better than a randomly chosen sequence. These ratios were averaged over all possible choices of the randomly chosen sequence and over the fifty trials, to yield the mean ratio score for a particular year and passage condition. A mean ratio score<1 indicates that the LBI algorithm performs better than random chance.

### 2.5 Statistical Analysis and Data Availability

Raw influenza sequences used in this analysis are available for download from GISAID (https://github.com/wilkelab/influenza_H3N2_passaging/tree/master/scripts.) using the parameters “North America”,“H3N2”, “1976 -2015”. Acknowledgements for sequences used in this study are available in SupplementaryFile 11. The complete, processed dataset used in our statistical analysis is available in Supplementary Data 10, including protein and gene numbering, computed evolutionary rates, relative solvent accessibility for the hemagglutinin trimer, and site-wise distance to protein site224. Relative solvent accessibility of the hemagglutinin trimer was taken from Meyer and Wilke (2015). Site-wise Euclidean distances between all amino acids in the HA structure PDBID:2YP7 were recalculated from structural coordinates using the script distances.py from:
https://github.com/wilkelab/influenzaH3N2passaging/tree/master/scripts. Statistical analysis was performed using R (Ihaka and Gentleman, 1996), and all graph figures were drawn with the R package ggplot2 (Wickham, 2009). Throughout this work, * denotes a significance of 0.01≤*P*<0.05, ** denotesa significance of 0.001 ≤ *P*<0.01, and *** denotes a significance of *P*<0.001.

Linear models between site-wise *dN/dS* and RSA or inverse distance were fit using the lm() function in R. Correlations were calculated using the R function cor() andsignificancedetermined using cor.test().Our entire analysis pipeline, instructions for running analyses, and raw data (except initial sequence data per the GISAID user agreement) are available at the following Github project repository:
https://github.com/wilkelab/influenza_H3N2_passaging.

## 3 RESULTS

Many influenza-virus samples collected from patients are first passaged through one or more culturing systems (Table 1) prior to PCR amplification and sequencing (Figure 1A). Samples may be passaged either once or serially (Table 1), even though a single passage is generally sufficient to obtain adequate amounts of viral DNA for sequencing. Reconstructed trees of influenza evolution contain a mixture of passage histories at their tips (Figure 1B). During passaging, influenza genomes accumulate adaptive mutations, and the effect of these mutations on evolutionary analyses of influenza sequences is not well understood.

**Figure 1.**
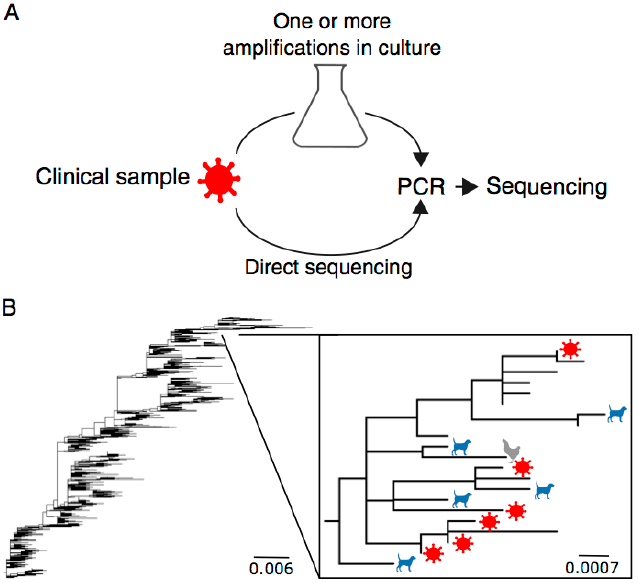
Schematic of influenza A virus sequence collection and analysis. (A) Typical processing steps of influenza A virus clinical isolates. Virus collected from patients may be passaged a single time or multiple times prior to PCR amplification and sequencing in a variety of different environments (Ex. canine cell culture, monkey cell culture, egg amniotes). However, some clinical virus is not passaged and is sequenced directly. (B) Phylogenetic tree of H3N2 HA sequences from the 2005-2015 seasons.The inset shows a small clade of sequences from the 2006/2007 season, with colored dots representing sequences with passage annotations(red virion: unpassaged, blue dog: canine cell culture, gray hen: egg amniote, unlabeled: missing or unclear passage history annotation).

### 3.1 Site-wise Evolutionary Rate Patterns Differ Between Passage Groups

To quantify any evolutionary signal that may be introduced by passaging, we assembled, from the GISAID database (Bogner et al., 2006), a set of North American human influenza H3N2 hemagglutinin sequences collected between 2005 and 2015.We initially sorted these sequences into groups by their passage history: (1) unpassaged, (2) egg-passaged, (3) generic cell-passaged, and (4) monkey cell-passaged (Table 1). To assess evolutionary variation at individual sites, we calculated site-specific *dN/dS* (Echave et al., 2016), using Single Likelihood Ancestor Counting (SLAC). Specifically, we calculated one-rate *dN/dS* estimates, i.e., site-specific *dN* values normalized by a global *dS* value (see Methods for details).In addition to considering groups of sequences with specific passage histories, we also calculated *dN/dS* values by pooling all sequences into one combined analysis. This pooled group corresponds to a typical influenza evolutionary analysis for which passage history has not been accounted.

We first correlated the site-wise *dN/dS* values we obtained for virus sequences derived from different passage histories. If passage history did not matter, then the*dN/dS* values obtained from different sources should have correlated strongly with each other, with *r* approaching 1. Instead, we found that correlation coefficients ranged from 0.68 to 0.87, depending on which specific comparison we made (Figure 2A). In this analysis, and throughout this work, we down-sampled alignments to the smallest number of sequences available for any of the conditions compared, to keep the samples as comparable as possible overall. The analysis of Figure 2 used *n*=917 randomly drawn sequences for each condition. Unpassaged*dN/dS*correlated more strongly with cell and pooled *dN/dS*> (correlations of 0.77 and 0.79, respectively) than with monkey-cell *dN/dS* (0.68J. Note that the *dN/dS* values from the pooled group, which corresponds to a typical dataset used in a phylogenetic analysis of influenza, more closely correlated with the *dN/dS* values from the generic cell group (r=0.87) than from the unpassaged group (r=0.79). Egg-derived sequences were excluded from this analysis due to low sequence numbers (n=79), however evolutionary rates from this condition correlated particularly poorly with those of random draws of 79 unpassaged sequences(Supplementary Figure 1). This result is consistent with previous conclusions (Bush et al., 2000; Gatherer, 2010; Suzuki, 2006) that egg-derived sequences show specific adaptations notfound otherwise in influenza sequences.

**Figure. 2.**
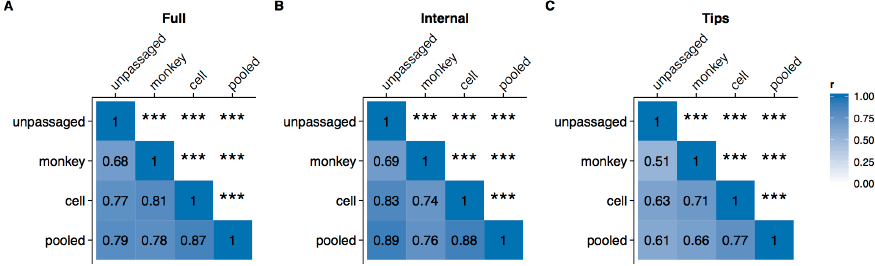
Comparison of sitewise *dN/dS* values among sequences with differing passage histories. Pearson correlations between sitewise *dN/dS* values for HA sequences derived from passaged and unpassaged influenza virus collected between 2005 and 2015 (downsampled to *n* = 917 in all groups). Correlations were calculated separately for *dN/dS* estimated from complete trees (A), internal branches only (B), and tip branches only (C). Asterisks denote significance of correlations (*0.01 ≤ P < 0.05,*0.001 ≤ P < 0.01, ***P < 0.001). Data used to generate this figure are available in Supplementary Data 1.

Because the common ancestor of any two passaged influenza viruses is a virus that replicated in humans, we expected that any adaptations introduced during passaging would not extend into the internal branches of a reconstructed tree. Therefore, we additionally subdivided phylogenetic trees into internal branches and tip branches, and calculated site-specific *dN/dS* values separately for these two sets of branches. In fact, Bush et al. (2000) recommended the use of internal branches to reduce variation seen between egg and cell culture-passaged virus. As expected, we found that when *dN/dS* calculations were restricted to the internal branches, the correlations between the passage groups increased overall (Figure 2B), even though distinct differences between the passage groups remained. Conversely, when we only considered tip branches, correlations among most groups were relatively low (Figure 2C), with the exception of cell-passaged sequences compared to the pooled sequences. This finding emphasizes once again that the pooled sample is most similar to the cell-passaged sample. We conclude that different passaging histories leave distinct, evolutionary signatures of adaptation to the passaging environment.

In aggregate, these results show that both generic-cell-passaged sequences and monkey-cell-passaged sequences yield different site-wise *dN/dS* patterns relative to unpassaged sequences (Fig. 2A-C), with *dN/dS* values derived from monkey-cell-passaged sequences being the least similar to *dN/dS* from unpassaged sequences (Fig. 2A-C). The pooled group of sequences, which corresponds to a typical dataset used in evolutionary analyses of influenza virus, describes evolutionary rates of specifically cell-passaged virus and poorly matches evolutionary rates of unpassaged virus.

**Figure. 3.**
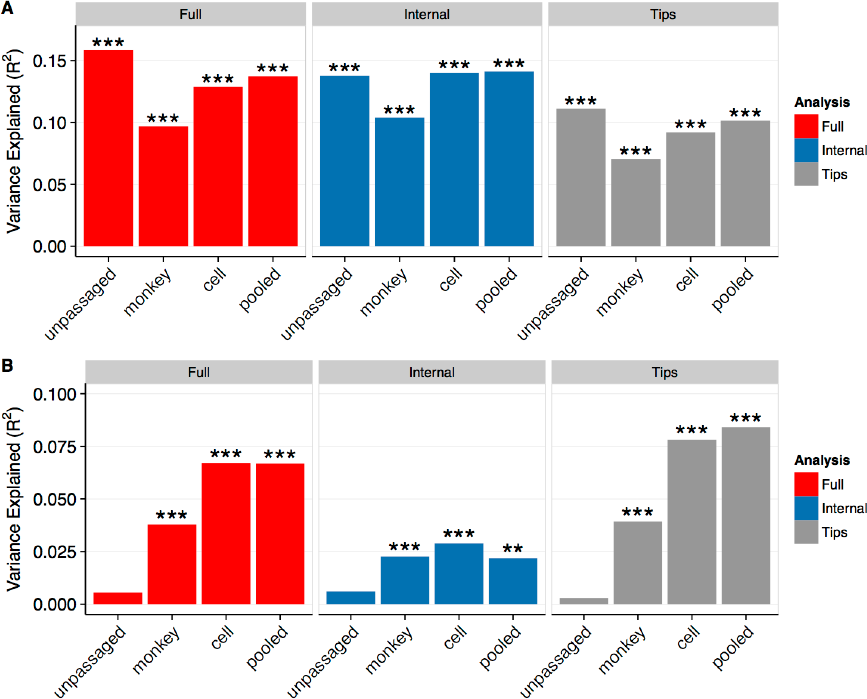
Percent variance in *dN/dS* explained by relative solvent accessibility (A)and by inverse distance to protein site 224 (B). (A) Relative solvent accessibility (RSA) explains ~10%.16% of the variation in *dN/dS* for all sequences. (B) Inverse distance to site 224 explains ~7% of the variation in *dN/dS* for cell-passaged sequences and for all sequences (pooled), however it explains virtually no variation for unpassaged sequences. Asterisks denote significance of correlations (*0.01 ≤ P < 0.05, ***0.001 ≤ P < 0.01, ***P < 0.001). Data used to generate this figure are available in Supplementary Data 1.

### 3.2 Adaptations to Cell and Monkey-cell Passage Display Characteristic Patterns of Site Variation

We next asked whether adaptations to passage history were located in specific regions of the hemagglutinin (HA) protein. To address this question, we employed the geometric model ofHA evolution we recently introduced (Meyer and Wilke, 2015), where structural measurements explain variation in *dN/dS*. For H3N2 HA, this model explains over 30% of thevariation in *dN/dS* using two simple physical measures, the relative solventaccessibility (RSA) of individual residues in the structure (Tien et al., 2013) and the inverse linear distance in 3D space from each residue to protein site 224 in the hemagglutinin monomer. Notably, the geometric model was previously applied to a pooled sequence set including sequences of various passaging histories. To what extent it carries over to sequences with specific passaging histories is not known.

We first considered the correlation between *dN/dS* and RSA (Figure 3A). We found that for all passage groups, R^2^ values ranged from 0.10 to 0.16 in the full tree, consistent with our earlier work (Meyer and Wilke, 2015). The high congruence among R^2^ values for internal branches and all branches suggests that RSA imposes a pervasive selection pressure on HA, independent of passaging adaptations. Thus, RSA represents a useful structural measure of a persistent effect of*dN/dS* with stronger correlations in the full tree and internal branches than in tip branches.

Next we considered the correlation between *dN/dS* and the inverse distance to site 224 to each site in the HA structure (Figure 3B). In contrast to RSA, correlations here were systematically higher in tip branches, suggesting a recent adaptive signal. We found virtually no correlation for unpassaged sequences, while a low correlation existed for monkey-cell cultured sequences and a higher correlation for cell-passaged and pooled sequences. To confirm that differences between pooled and unpassaged correlations were not simply due to variation from random sampling, we created a null distribution of correlations from 200 random draws of pooled sequences. The correlation for unpassaged sequences was significantly lower than it was for pooled sequences (z=— 4.22, *p*=1.2 × 10^—5^, Supplementary Figure 2). Correlations from pooled sequences mirrored cell-culture correlations and persisted through internal branches. Thus, the correlation of *dN/dS* with the inverse distance to site 224 seems to be primarily an artifact of cell passage, even though its effect can be seen along internal branches as well. This cell-specific signal dominates the pooled dataset. Further, this cell-specific signal is partially attenuated along internal branches and amplified along tip branches, as we would expect from a signal caused by recent host-specific adaptation. Even though this signal is a true predictor of influenza evolutionary rates for virus grown in cell culture, it does not transfer to unpassaged sequences and therefore has no relevance for the circulating virus. This finding serves as a strong demonstration of passage history as a confounder in analysis of hemagglutinin evolution, not just for egg passage as previously demonstrated, but also for cell and monkey-cell passage.

Surprisingly, the correlation we found here between *dN/dS* and inverse distance to site 224 for pooled sequences *(R^2^*=0.067) was less than half of the value previously reported (Meyer and Wilke, 2015) (Fig. 3B). However, using a dataset of sequences more temporally matched tothe previouslypublished analysis (2005-2014 instead of 2005-2015), we recovered the earlier higher correlation. This finding suggests that there is some feature in the additional 2015 sequences that changes the pooled data's relationship with inverse distance to site 224. In 2015, unpassaged and SIAT1 sequences each doubled in number compared to 2014, while the number of non-SIAT1 cell cultured sequences dropped dramatically (Table 1). SIAT1, an MDCK cell line which overexpresses human-like 6-linked sialic acids over native 3-linked sialic acids (Matrosovich et al., 2003) has higher sequence fidelity than unmodified MDCK (Hamamoto et al., 2013). Therefore, we next investigated whether the drop in correlation from 2014 to 2015 could be attributed to the recent reduction in cell culture using non-SIAT1 cells.

### 3.3 Adaptation to Passage in SIAT1 Cells is Weak or Absent

In the preceding analyses, we lumped all cell cultures except monkey cells into the same category. However, there are more subtle distinctions in cell passaging systems, and they can exert differential selective pressures on human adapted virus (Hamamoto et al., 2013; Oh et al., 2008). As our generic cell culture group was composed of a mixture of wild type MDCK,SIAT1, and unspecified cell cultures, we next investigated whether any one culture type was the source of the high cell-culture signal seen in Figure 3B.

SIAT1 is currently the dominant system for passaging of influenza virus in North America with approximately half of the 2015 influenza sequences currently available from GISAID deriving from serial passaging through SIAT1 cells. Experimental analysis of SIAT1 demonstrates improved sequence fidelity and reduced positive selection over unmodified MDCK cell culture (Hamamoto et al., 2013; Oh et al., 2008). We sought to determine if the apparently cell-culture-specific correlation of site-wise evolutionary rates and inverse distance to site 224 extended to SIAT1 cell culture. To compare cell-culture varieties, we created sample-size matched groups of non-SIAT1 cell culture, SIAT1 cell culture, and unpassaged sequences collected between 2005 and 2015 (*n*=1046), excluding sequences that had been passaged through both a non-SIAT1 and a SIAT1 cell culture.

All groups showed similar correlations between *dN/dS* and RSA, regardless of whether *dN/dS* was calculated for the entire tree, for internal branches only, or for tip branches only (Figure 4A). By contrast, inverse distance to site 224 uniquely correlated with *dN/dS* from non-SIAT1-cultured virus (Figure 4B). This effect was strongest along tip branches *(R^2^*=0.139), but it was almost as strong along the entire tree *(R^2^*=0.129). The correlation was reduced, though still significant, among internal branches (*R^2^*=0.075). Thus, we conclude that the correlation between *dN/dS* and the inverse distance to site 224 represents a unique signal of adaptation to passaging in non-SIAT1 cells. In other words, a non-SIAT1-specific signal can completely dominate all signals ofpositive adaptation when a datasetcontains a sufficiently high number of sequences passaged in non-SIAT1 cells. In our analysis(Figure 3B), the high correlation of non-SIAT1 cell *dN/dS* with inverse distance to site 224 is suppressed in the pooled condition because the number of unpassaged and SIAT1-passaged sequences grew substantially in 2015. This difference in sample composition explains the lower than expected correlations in Figure3B for pooled *dN/dS*.

**Figure. 4.**
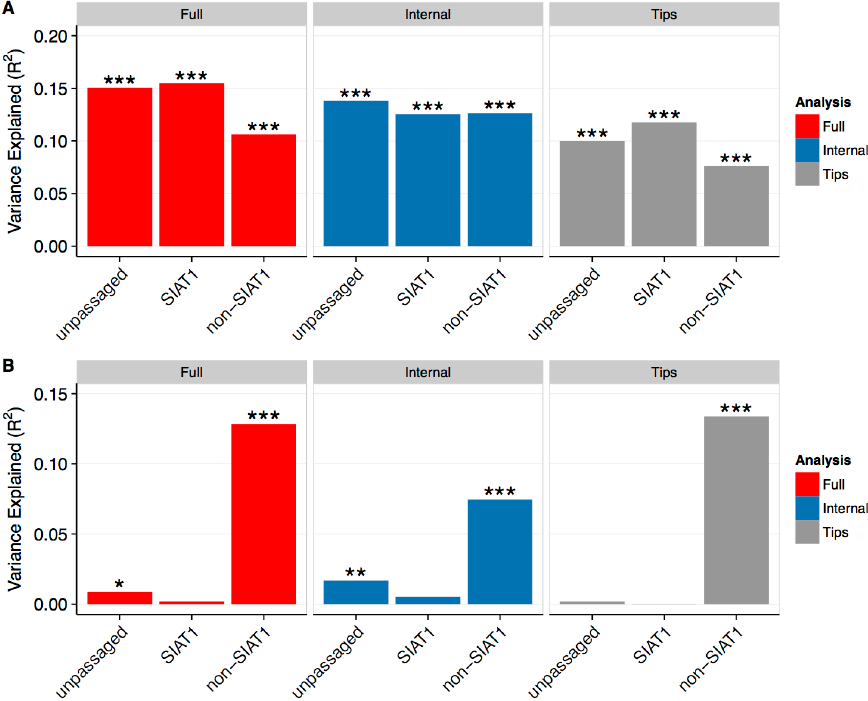
Virus passaged in non-SIAT1 cells carries unique adaptations not present in unpassaged or SIAT1-passaged virus. (A) The correlation between *dN/dS* and RSA is weakened for virus passaged in non-SIAT1 cells. (B) The correlation between *dN/dS* and inverse distance to site 224, representing a positive-selection hotspot in the vicinity of that site, is only present in virus passaged in non-SIAT1 cells. Asterisks denote significance levels (*0.01 ≤ P <0.05, **0.001 ≤ P < 0.01, ***P < 0.001). Sequences analyzed were collected between 2005 and 2015. Alignments were randomly down-sampled to yield identical numbers of sequences in each alignment (n = 1046). Data used to generate this figure are available in Supplementary Data 4.

As these three conditions were somewhat temporally separated (most non-SIAT1 cell culture sequences were pre-2015, and most unpassaged and SIAT1 culture sequences were post-2014), we controlled for season-to-season variation by drawing 249 sequences from each group from 2014 We again considered site-wise *dN/dS* correlations among passaging groups,and we found that overall, unpassaged and SIAT1-passaged sequences appeared the most similar (Supplementary Figure 3A-C).

### 3.4 Signals of Passaging Adaptation Accumulate with Additional Rounds of Passaging in Non-SIAT1 Cells

Having identified non-SIAT1 cell culture as the source of the contaminating signal in analyses of inverse distance to site 224, we next investigated the source of this signal at the single amino acid level. We expected that a signal of adaptation to a passaging system would strengthen with additional exposure to that system. Thus, we compared the magnitude of the site-wise *dN/dS* values in sequences that had never been passaged, had been passaged once, passaged twice, or passaged three to five times (Figure 5A). For this analysis, we only considered passage in non-SIAT1 cells. This analysis revealed distinct regions of increasing positive selection along the hemagglutinin molecule (arrows in Figure 5A) and a strong relationship between the magnitude of these signals and the number of times influenza viruses were passaged. Further, we found an overall increase in *dN/dS* with increased numbers of passages in non-SIAT1 cells (Figure 5B). The strongly selected sites 221 and 225 are adjacent to site 224, explaining the specific relationship between *dN/dS* calculated from non-SIAT1 sequences and the inverse distance in 3D space to this site. The correlation between *dN/dS* and inverse distance increased in strength with increasing numbers of passages (Figure 5C), even though it was observable after a single passage in non-SIAT1 cells. Mapping the raw *dN/dS* values onto the hemagglutinin structure showed how specific sites light up as passage numbers increase (Figure 5D).

**Figure. 5.**
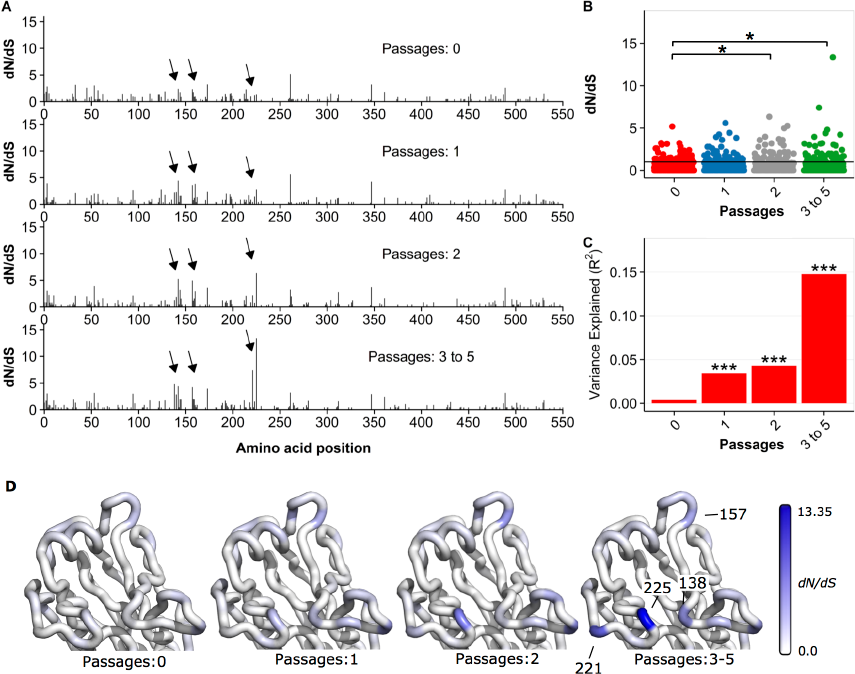
Accumulation of passaging artifacts with increasing numbers of serial passages in non-SIAT1 cell culture. (A) Sitewise *dN/dS* values for virus which was not passaged, passaged once, passaged twice, and passaged three to five times in non-SIAT1 cell culture (*n* = 304for each group). Arrows highlight regions of increased *dN/dS* in passaged virus. Notably, *dN/dS* inflation is increased with increasing rounds of passaging. (B) *dN/dS* values vs. number of passages. *dN/dS* values are significantly elevated after two or more passages in non-SIAT1 cell culture, relative to unpassaged virus (paired t test). Asterisks denote significance levels (*0.01 ≤ P < 0.05, **0.001 ≤ P < 0.01, ***P < 0.001). (C) The correlation between *dN/dS* and inverse distance to site 224 increases with the number of passages. (D) Mapping *dN/dS* values onto the hemagglutinin head structure demonstrates the accumulation of 911 passage adaptations with increasing rounds of passages. Labeled sites correspond to regions denoted with arrows in (A). Data used to generate this figure are available in Supplementary Data 6.

### 3.5 Evolutionary Variation in Sequences from Unpassaged Virus Predicts Regions Involved in Antigenic Escape

Our preceding analyses might suggest that the inverse distance metric for describing regions of selection only captures effects of adaptation to non-SIAT1 cell culture. However, this is not necessarily the case. Importantly, inverse distance needs to be calculated relative to a specific reference point. Site 224 was previously used as the reference point because it yielded the highest correlation for the dataset analyzed (Meyer 2015 and Wilke, 2015). For a different dataset, one that doesn’t carry the signal of adaptation to non-SIAT1 cell culture, a different reference point may be more appropriate.

We thus repeated the inverse distance analysis of Meyer 2015 and Wilke (2015) for a size-matched sample of 1046 sequences from non-SIAT1, pooled, SIAT1, and unpassaged virus collected between 2005 and 2015 (Figure 6). In brief, for each possible reference site in the hemagglutinin structure, we measured the inverse distance in 3D space from that site to every other site in the structure (see Methods for details). We then correlated the inverse distances with the *dN/dS* values at each site, resulting in one correlation coefficient per reference site. Finally, we mapped these correlation coefficients onto the HA structure, coloring each reference site by its associated correlation coefficient. If inverse distances measured from a particular reference amino acid have higher correlation with the site-wise *dN/dS* values, then this reference site will appear highlighted on the structure.

**Figure. 6.**
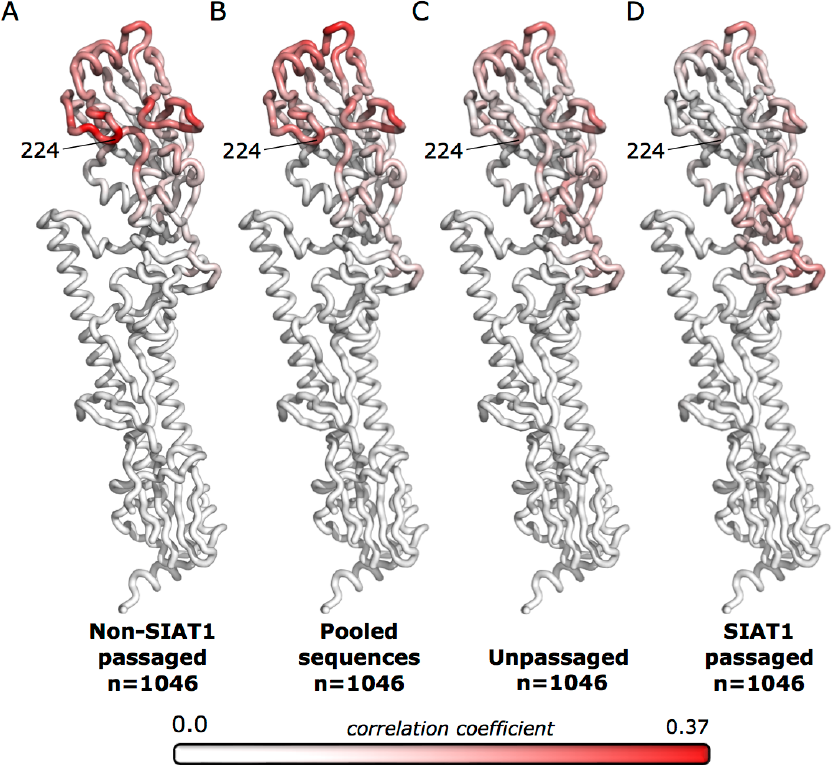
Correlations of *dN/dS* with inverse distances, mapped onto the hemagglutinin structure for non-SIAT1-passaged, pooled, unpassaged, and SIAT1 passaged sequences. The correlation between *dN/dS* and inverse distance for each reference site was mapped onto the hemagglutinin structure for (A) non-SIAT1 sequences, (B) pooled sequences, (C) unpassaged sequences, and (D) SIAT1 passaged sequences. Sequences analyzed were collected between 2005 and 2015. Alignments were randomly down-sampled to yield identical numbers of sequences in 45 each alignment (*n* = 1046). Red coloring represents positive correlations, while white represents zero or negative correlations. The four conditions group into two distinct correlation patterns, non-SIAT1/pooled and unpassaged/SIAT1. In particular, the loop containing site 224 lights up strongly for non-SIAT1 and pooled sequences but not for unpassaged and SIAT1 sequences. Data used to generate this figure are available in Supplementary Data 7.

For non-SIAT1-passaged and pooled virus, this analysis recovered the finding of Meyer and Wilke (2015) that the loop containing site 224 appeared strongly highlighted (Figure 7A). However, this signal was entirely absent in unpassaged and SIAT1 passaged virus (Figure 7B), with no sites in that loop working well as a reference point. These results suggested that this loop was specifically involved in adaptation of hemagglutinin to non-SIAT1 cell culture, explaining the non-SIAT1-specific signal shown in Figure 4A. Globally, the pattern of correlations from pooled sequences strongly resembled the non-SIAT1 pattern, in contrast to the resemblance of SIAT1 to unpassaged. Thus, the inverse distance metric is useful for differentiating regions of selection particular to different experimental groups.

**Figure. 7.**
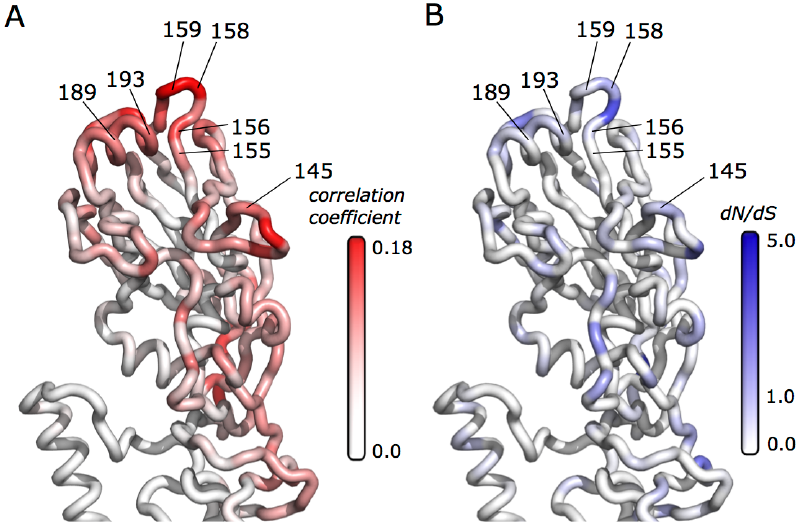
Unpassaged sequences allow recovery of antigenic regions from positive-selectionanalysis. For each site, the correlation between *dN/dS* and invers distance (A) or *dN/dS* directly (B) were mapped onto the hemagglutinin structure, for *dN/dS* derived from unpassaged sequences collected between 2005 and 2015 (*n* = 1703). Red coloring represents higher correlation; blue coloring represents higher *dN/dS*. Highlighted regions contain residues (labeled withprotein site number) which experimentally determined to cause antigenic change by Koel et al., 2013. Correlations and *dN/dS* for antigenic residues are given in Table 2. Data used to generate this figure are available in Supplementary Data 7.

Therefore, we next asked what residual patterns of positive selection remained once the adaptation to non-SIAT1 cells was removed. Even though site-wise correlations are relatively low for unpassaged virus compared to the ones observed for non-SIAT1-passaged virus, we could still recover relevant patterns of HA adaptation after rescaling our coloring. In particular, we found that sites opposite to the loop-containing site 224 lit up in our analysis of unpassaged sequences (Figure 7A). Sites in this region are known to be involved in antigenic escape. In fact, many of the highlighted regions contain amino acid positions where substitution led to antigenic change (Table 2). We found a similar pattern of concordance with antigenic sites when mapping *dN/dS* values directly onto the structure (Figure 7B). The inverse-distance correlations, however, performed better at identifying antigenic residues than did raw *dN/dS* values. When considering the 90^th^ percentile (top 10% highest scored sites) by either metric, the inverse-distance correlations recovered 5 of 7 sites while *dN/dS* alone recovered only 1 of 7 sites (Table 2). Additionally, while several sites involved in antigenic change had very low *dN/dS*, all had inverse-distance correlations above the 86^th^ percentile.

**Table 2.** Evolutionary rates and inverse distance correlations 850 of residuesresponsible for antigenic change. For each site, we determined *dN/dS* and the correlation between *dN/dS* and inverse distance for unpassaged sequences collected between 2005 and 2015 (*n* = 1703). 5/7residues linked to antigenic changes have inverse-distance correlations above the 90th percentile, while only 1/7 have *dN/dS* values above the 90th percentile. Sites were experimentallydetermined by Koel et al., (2013).

| Site | | Raw $dN/dS$ | | Inv.-dist. correlation | |
| --- | --- | --- | --- | --- | --- |
| Gene | Protein | $dN/dS$ | percentile | $r$ | percentile |
| 161 | 145 | 0.672 | 0.823 | 0.082 | 0.883 |
| 171 | 155 | 0 | 0 | 0.077 | 0.867 |
| 172 | 156 | 0.672 | 0.832 | 0.1317 | 0.971 |
| 174 | 158 | 1.36 | 0.958 | 0.1797 | 0.996 |
| 175 | 159 | 0.49 | 0.75 | 0.1837 | 1 |
| 205 | 189 | 0.474 | 0.763 | 0.0887 | 0.905 |
| 209 | 193 | 0.672 | 0.845 | 0.098 | 0.936 |

### 3.6 Passaging Artifacts Extend Deep into Reconstructed Trees

Passaging adaptations could reasonably be expected to only affect peripheral clades of influenza virus evolutionary trees, as they represent recent signals of adaptation that should not penetrate far into the tree or significantly affect tree structure (Kryazhimskiy and Plotkin, 2008; Strelkowa and Lässig, 2012). Surprisingly, however, we found signals of passage adaptations in season-to-season fixed mutations and branch density.

To capture mutations that became fixed across seasons, we calculated *dN/dS* along the trunks of trees constructed from sequences of difference passage histories (Figure 8A). To time-calibrate our trunk, we limited this analysis to passage types that had sequences at least in 2005 and in 2015, which excludes SIAT1 sequences. Trunk *dN/dS* measures season-to-season adaptation and might be expected to be robust to the effects of passaging. However, recurring adaptations to passaging conditions as samples were processed across seasons could falsely appear as trunk mutations.

**Figure. 8.**
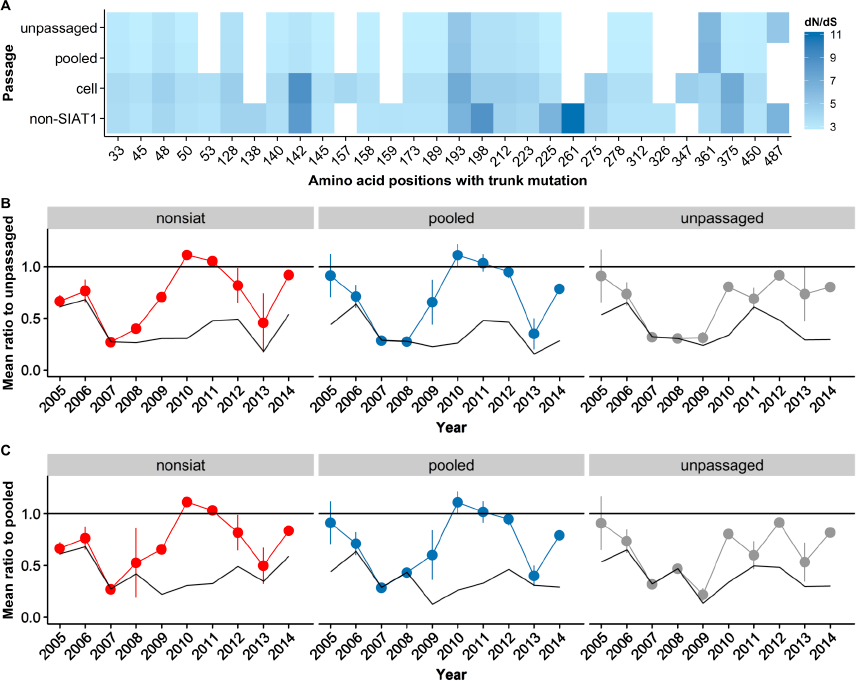
Passage artifacts affect trunk *dN/dS* and topology-based predictions. (A) Sitewise trunk *dN/dS* values for passage groups (*n* = 1703). Only sites with at least one non-synonymous mutation along the trunk were included. Many sites appear underpositive selectiononly in the trunk reconstructed from passaged sequences, e.g. 159, 261, 275. (B, C) Prediction of future dominant clade by Local Branching Index (LBI) depends on passaging history. LBIwas calculated for trees derived from non-SIAT1, pooled, and unpassaged sequences using a maximum of 100 sequences per condition. The mean ratio estimates the quality of the prediction(lower is better), and values < 1 indicate the prediction performs better than rando.Error bars indicate the standard 48deviation of the mean, estimated from resampling (Methods.948 The solid black lines represent the best possible prediction in each year. Predictions were evaluated relative to the following year’s ancestrally reconstructed root sequence obtained from (A) unpassaged and (B) pooled sequences. Non-SIAT1, pooled, and unpassaged sequences have divergent prediction quality across years, and on average, unpassaged sequences seem to perform better than passaged or pooled sequences. (The distances between dots and the solid black lines are, on average, the smallest for predictions derived from unpassaged sequences.) Data used to generate this figure are available in Supplementary Dataset 8 (A) and Supplementary Dataset 9 (B and C).

For pooled sequences, trunk *dN/dS* appeared to be generally free of artifacts, resembling the trunk *dN/dS* of unpassaged sequences (Figure 8A). In contrast, for datasets composed entirely of passaged sequences we found artifacts extending into the trunk. When trees were constructed from only cell-passaged sequences or only non-SIAT1 sequences, we observed a general inflation in *dN/dS* as well as several spurious sites of high *dN/dS* that do not occur in the unpassaged condition. Together, this result shows that the relative proportion of passaged to unpassaged sequences in a sample matters; when sequences with passaging artifacts are overrepresented compared to unpassaged sequences, there is a risk that a spurious signal will be found in the trunk. For example, a trunk *dN/dS* analysis of mainly non-SIAT1 sequences would direct attention to site 261, even though this site does not appear to be positively selected on the unpassaged tree trunk. This analysis demonstrates the ability of sequences containing major passaging artifacts to confound both deep and peripheral analyses of influenza virus evolution.

We next investigated the effect of passaging on a tree-topology based metric, Local Branching Index (LBI) (Neher et al., 2014). Notably, this metric is entirely independent of *dN/dS* values. The LBI algorithm uses the degree of local branching around a terminal node to predict sequences similar to progenitors of the following season’s strain. As a read-out of the algorithm’s performance,we calculated the mean ratio score, as described (Neher et al., 2014) (see also Methods). Mean ratio scores below 1 indicate that the algorithm performs better than random chance. The lowest possible mean ratio score possible corresponds to the mean ratio score of the sequence with the lowest hamming distance to the following year’s progenitor (i.e., the theoretical best possible prediction from sequences in a condition).

We saw clear differences in accuracy of predictions made using trees composed of non-SIAT1,pooled, or unpassaged sequences (Figure 8B, C). We could not examine SIAT1 patterns, as these sequences were not consistently available across seasons until 2013. As a general trend, predictions from unpassaged sequences seemed to be more accurate (both less likely to exceed 1 and more likely to be closer to the best possible prediction) than predictions from either passaged orpooled sequences.

## 4 DISCUSSION

We find that serial passaging of influenza virus introduces a measurable signal of adaptation into the evolutionary analysis of natural influenza sequences. There are unique, characteristic patterns of adaptation to egg passage, monkey cell passage, and non-SIAT1 cell passage. Monkey-cell-derived sequences show different molecule-wide evolutionary rate patterns. Non-SIAT1 cell-derived sequences instead display a hotspot of positive selection in a loop underneath the sialic-acid binding region. This hotspot has been previously noted (Meyer and Wilke, 2015) but no explanation for its origin was available. Additional passages in non-SIAT1 cell strengthen this artifact. Further, we find that virus passaged in SIAT1 cells seems to accumulate only minor passaging artifacts. Throughout our analyses, we find limited utility in subdividing phylogenetic trees to internal and terminal branches. While signals of passage adaptation are consistently elevated along terminal branches and attenuated along internal branches, evolutionary rates along internal branches remain confounded by passaging artifacts. Additionally, passage adaptation can resemble fixed season-to-season mutation along trunk branches and alter topology-based predictions of sequence fitness. Finally, we can accurately recover the experimentally determined antigenic regions of hemagglutinin from evolutionary-rate analysis by using a dataset consisting of only unpassaged viral sequences.

Previous studies (Bush et al., 2001; Suzuki, 2006) suggest the use of internal branches to alleviate passage adaptations. However, we find here that this strategy is insufficient, because the evolutionary signal of passage adaptations can often be detected along internal branches. This finding may seem counterintuitive, as internal nodes should exclusively represent human-adapted virus. We suggest that passaging adaptations in internal branches may be homoplasies caused by convergent evolution; if different clinical isolates converge onto the same adaptive mutations under passaging, then these mutations may incorrectly be placed along internal branches under phylogenetic tree reconstruction. Additionally, although the use of only internal branches removes some differences between the passage groups, the exclusion of terminal sequences can obscure recent natural adaptations and thus obscure actual sites under positive selection. Therefore, analysis of internal branches is not only insufficient for eliminating artifacts from passaging adaptations but also suboptimal for detecting positive selection in seasonal H3N2 influenza.

The safest route to avoid passaging artifacts is to limit sequence datasets to only unpassaged virus, although this approach limits sequence numbers. The human-like 6-linked sialic acids in SIAT1 (Matrosovich et al., 2003) greatly reduce observed cell culture-specific adaptations, particularly in the loop of hemagglutinin which contains site 224. This lack of selection concords with multiple experiments finding low levels of adaptation in this cell line(Hamamoto et al., 2013; Oh et al., 2008). As our analysis only detects minor differences between unpassaged and SIAT1 passaged virus, we posit that this passage condition is an acceptable substitute for unpassaged clinical samples. Even so, our findings do not preclude the existence of SIAT1-specific adaptations that may confound specific analyses.

Over half of the passaged hemagglutinin sequences in the GISAID database from 2005-2015 were passaged more than once. Multiple passages cause increasing accumulation of passaging artifacts in non-SIAT1 cells, and we predict that any yet unknown passaging effects would accumulate similarly. However, even a single passage in non-SIAT1 cells introduces noticeable artifacts of adaptation. Influenza virus is often passaged multiple times to improve viral titers for hemagglutination inhibition assays, and thus, we expect that multiply passaged viruses will continue to be deposited for the foreseeable future. We recommend that such viruses be used with care when studying the evolutionary dynamics of influenza strains circulating inthe human population.

Although the majority of the sequences from the year 2015 are SIAT1-passaged or unpassaged, several hundred sequences from that year derive from monkey cell culture. The use of monkey cell culture surged in 2014 and 2015 compared to previous years. We recommend that these recently collected sequences be excluded from influenza rate analysis, in favor of the majority of unpassaged and SIAT1-passaged sequences. As passaging is a useful and cost effective method for amplification of clinically collected virus, unpassaged viral sequences are unlikely to completely dominate influenza sequence databases in the near future. However, new human epithelial cell culture systems for influenza passaging (Ilyushina et al., 2012) could soon provide an ideal system that both amplifies virus and protects it from non-human selective pressures.

Passage history should routinely be considered as a potential confounding variable in future analyses of influenza evolutionary rates. Future studies should be checked against unpassaged samples to ensure that conclusions are not based on adaptation to non-human hosts. We recommend the exclusion of viral sequences that derive from serial passage in egg amniotes, monkey kidney cell culture, and any unspecified cell culture. Prior work that did not consider passaging history may be confounded by passaging adaptations, as occurred in our previouspublication (Meyer 2015 and Wilke, 2015). In that manuscript, we concluded that sites under positive selection differ from sites involved in immune escape. Here, we find that the origin ofthis positive selection is adaptation to the non-human passaging host, not immune escape in or adaptation to humans. In particular, we suggest that the evolutionary markers of influenzavirus determined in (Belanov et al., 2015) be reevaluated to ensure these sites are not artifacts of viral passaging. Similarly, many of the earlier studies (Bush et al., 1999; Meyer and Wilke, 2013, 2015; Pan 2011 and Deem, 2011; Shih et al., 2007; Suzuki, 2006, 2008; Tusche et al., 2012) performing site-specific evolutionary analysis ofhemagglutinin likely contain some conclusions that can be traced back to passagingartifacts.Additionally, even though passage artifacts do not appear to be sufficiently strong to affect clade-structure reconstruction (Bush et al., 2000), they do have the potential to cause artificially long branch lengths, due to *dN/dS* inflation, or misplaced branches, due to convergent evolution under passaging. We find samples composed of non-SIAT1 appear to behave differently than unpassaged samples under the Local Branching Index metric. Thus, future phylogenetic predictive models of influenza fitness and antigenicity, as in (Bedford et al., 2014; Łuksza and Lässig, 2014; Neher et al., 2014), should also be checked for robustness to passage-related signals. Finally, while it is beyond the scope of this work to investigate passage history effects in other viruses, we suspect that passage-derived artifacts could be a factor in their phylogenetic analyses as well. The use of datasets free of passage adaptations will likely bring computational predictions of influenza positive selection more in line with corresponding experimental results.

Sequences without passage annotations are inadequate for reliable evolutionary analysis of influenza virus. Yet, passage annotations are often completely missing from strain information, and, when present, are often inconsistent; there is currently no standardized language to represent number and type of serial passage. We note, however, that passage annotations from the 2015 season are greatly improved when compared to previous seasons. Several major influenza repositories, including the Influenza Research Database (Squires et al., 2012) and the NCBI Influenza Virus Resource (Bao et al., 2008), do not provide any passaging annotations at all. Additionally, passage history is not required for new sequence submissions to theNCBI Genbank (Benson et al., 2012). The EpiFlu database maintained by the Global Initiative for Sharing Avian Influenza Data (GISAID) (Bogner et al., 2006) and OpenFluDB (Liechti et al., 2010), however, stand apart by providing passage history annotations for the majority of their sequences. Of these, only the OpenFluDB repository allows filtering of sequences by passage history during data download. Our results demonstrate the strength of passaging artifacts in evolutionary analysis of influenza. The lack of a universal standard for annotation of viral passage histories and a universal standard for serial passage experimental conditions complicate the analysis and mitigation of passaging effects.

## Acknowledgements

We would like to thank Sebastian Maurer-Stroh for help with interpreting passaging annotations in GISAID. This work was supported in part by NIH grant no. R01GM 088344, DTRA grant no. HDTRA1-12-C-0007, and NSF Cooperative agreement no.DBI-0939454 (BEACON Center). The funders had no role in study design, data collection and analysis, decision to publish, or preparation of the manuscript

## DATA AVAILABILITY

Processed data are available as supplementary material. Sequence data are available from GISAID as detailed in Methods. All analysis code used to generate the processed data is available at: https://github.com/wilkelab/influenzH3N2passaging

## AUTHOR CONTRIBUTIONS

Conceived and designed the experiments: CDM COW. Wrote scripts and analytic tools: CDM AGM. Performed the experiments: CDM. Analyzed the data: CDM COW. Wrote the paper: CDM AGM COW.

## SUPPLEMENTARY FIGURES AND DATA

**Supplementary Figure 1.**
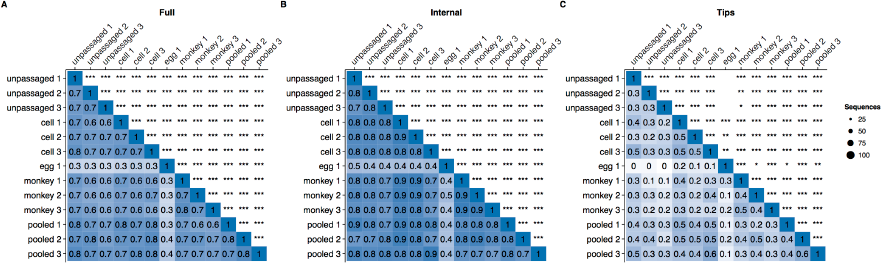
**Comparison of sitewise hemagglutinin *dN/dS* derived, from size-matched samples of sequences with various passage histories, including egg amniotes**. Pearson correlations between sitewise *dN/dS* values for HA sequences derived from passaged and unpassaged influenza virus collected between 2005 and 2015, randomly down-sampled to 79 sequences per passage group. (Since there were so few egg-derived sequences, each down-sampling was independently performed three times, resulting in replicates 1, 2, and 3 for each passage group.) Correlations were calculated separately for complete trees (A), internal branches only (B), and tip branches only (C). Asterisks denote significance of correlations (*0.01 ≤ P <969 0.05, **0.001 ≤ P < 0.01, ***P < 0.001). Data used to generate this figure are available in Supplementary Data 2.

**Supplementary Figure 2.**
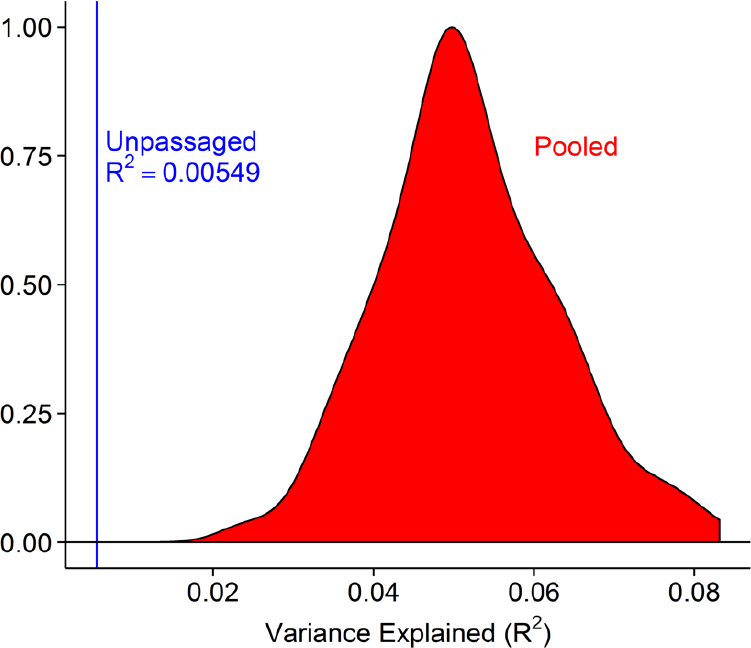
**Differences in correlation strength not due to variance from random sampling**. Percent variance in *dN/dS* explained by inverse distance to site 224 for unpassaged and pooled sequences (*n* = 917). For pooled sequences, sets of 917 sequences were randomly drawn 200 times, with replacement. For each set, we calculated *dN/dS* and correlated with inverse distance. The R^2^ for unpassaged sequences falls well outside the distribution of R^2^ values for pooled sequendes (*z* = −4.22, *p* = 1.2 × 10^−5^). Data used to generate this figure are available in Supplementary Data 3.

**Supplementary Figure 3.**
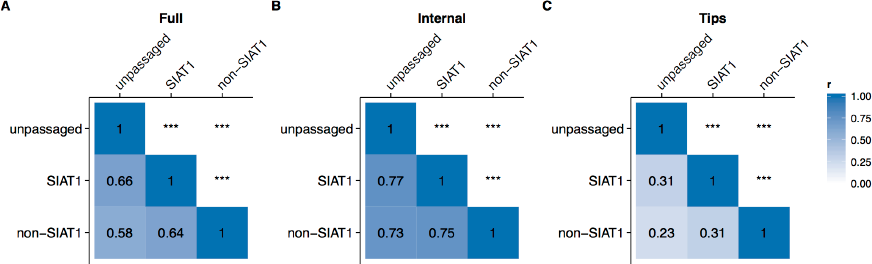
**Comparison of unpassaged, non-SIAT1-passaged, and SIAT1-passaged virus in 2014 only**. (A–C) Pearson correlations between sitewise *dN/dS* values for the three passage groups (*n* = 249), for complete trees, internal branches only, and tip branches only, respectively. Data used to generate this figure are available in Supplementary Data 5.

**Supplementary Figure 4.**
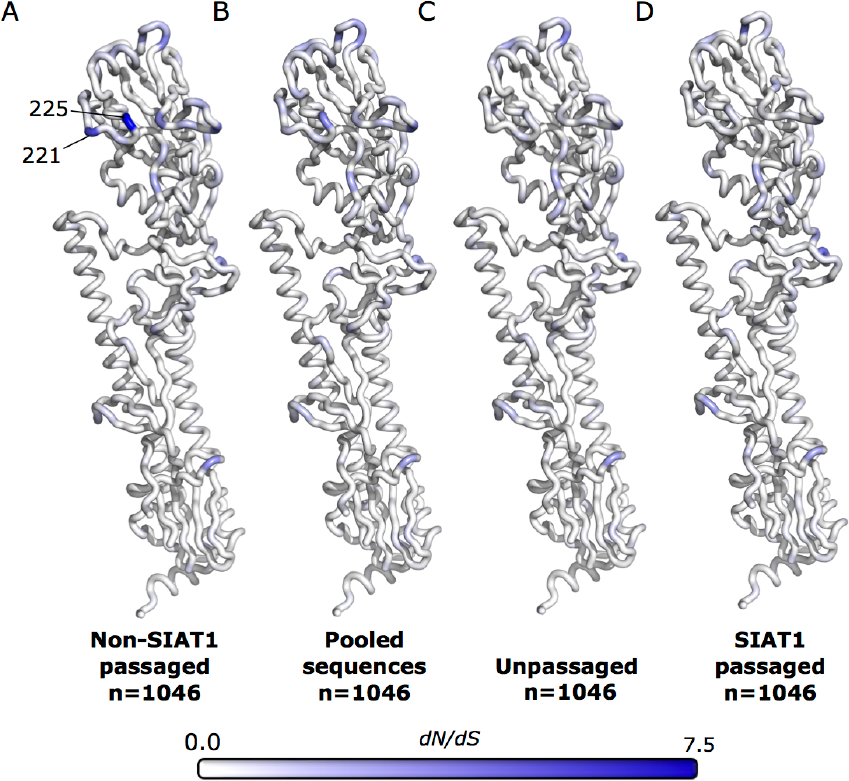
***dN/dS* mapped onto the hemagglutinin structure for non-SIAT1-passaged, pooled, unpassaged, and SIAT1 passaged sequences**. *dN/dS* was mapped onto the hemagglutinin structure for (A) non-SIAT1 sequences, (B) pooled sequences, (C) unpassaged sequences, and (D) SIAT1 passaged sequences collected between 2005 and 2015 (*n* = 1046). dN/dS from pooled sequences appears to be intermediate between dN/dS from non-SIAT1-passaged and unpassaged sequences, particularly at sites 221 and 225. Data used to generate this figure are available in Supplementary Data 7.

**Supplementary Data 1**. Data used to generate Figures 2 and 3. 997 This file includes 1)sitewise *dN/dS* values of random draws of 917 unpassaged, generic cell cultured, monkey cell cultured, and the pooled group sequences collected between 2005 and 2015, 2) protein and gene numbering, 3) PDB:2YP7 sequence, 4) relative solvent accessibilities of the hemagglutinin trimer, 5) linear distances to protein site 224.

**Supplementary Data 2**. Data used to generate Supplementary Figure 1. This file includes 1) sitewise *dN/dS* values of random draws of 97 unpassaged, generic cell cultured, egg cultured, monkey cell cultured,and the pooled group sequences collected between 2005 and 2015, 2) protein and gene numbering, 3) PDB:2YP7 sequence.

**Supplementary Data 3**. Data used to generate Figure Supplementary Figure 2. This file includes 1) sitewise *dN/dS* values of 500 random draws of 917 pooled sequences and one column of sitewise dN/dS from 917 unpassaged sequences collected between 2005 and 2015, 2) protein and gene numbering, 3) PDB:2YP7 sequence 4) linear distances to protein site 224.

**Supplementary Data 4**. Data used to generate Figure 4. This file includes 1) sitewise *dN/dS* values of random draws of 1046 unpassaged, SIAT1, and non-SIAT1 cell culture sequences collected between 2005 and 2015, 2) protein and gene numbering, 3) PDB:2YP7 sequence, 4) relative solvent accessibilities of the hemagglutinin trimer, and 5) linear distances to protein site 224.

**Supplementary Data 5**. Data used to generate Supplementary 1020 Figure 3. This file includes 1) sitewise *dN/dS* values of random draws of 249 unpassaged, SIAT1 cultured, and non-SIAT1 cell cultured sequences collected in 2014, 2) protein and gene numbering, 3) PDB:2YP7 sequence.

**Supplementary Data 6**. Data used to generate Figure 5. This file includes 1) sitewise *dN/dS* values of random draws of 304 unpassaged and 304 non-SIAT1 cell cultured sequences passaged once, twice, or 3-5 times collected between 2005 and 2015, 2) protein and gene numbering, 3) PDB:2YP7 sequence.

**Supplementary Data 7**. Data used to generate Figures 6, 7, and Supplementary Figure 4. This file includes 1) sitewise *dN/dS* values of random draws of 1703 unpassaged and non-SIAT1 cell cultured sequences collected between 2005 and 2015, 2) protein and gene numbering, 3) PDB:2YP7 sequence, 4) sitewise inverse distance correlations.

**Supplementary Data 8**. Data used to generate Figure 8A. This file includes 1) trunk sitewise *dN/dS* values of all available unpassaged, SIAT1 cultured, non-SIAT1 cell, generic cell culture, monkey cell cultured, and pooled sequences collected from 2005-2015, 2) protein and gene numbering, 3) PDB:2YP7 sequence.

**Supplementary Data 9**. Data used to generate Figure 8B and C. This file includes output of Local-Branching-Indexanalyses.

**Supplementary Data 10**. This file includes 1) all sitewise 1043 *dN/dS* values used to generate figures, 2) protein and gene numbering, 3) PDB:2YP7 sequence, 4) relative solvent accessibilities of the hemagglutinin trimer, and 5) linear distances to proteinsite 224.

**Supplementary Data 11**. GISAID acknowledgements for hemagglutinin sequences collected between 1968 and 2015.

